# Revisiting Parameter Estimation in Biological Networks: Influence of Symmetries

**DOI:** 10.1101/674739

**Authors:** Jithin K. Sreedharan, Krzysztof Turowski, Wojciech Szpankowski

## Abstract

Graph models often give us a deeper understanding of real-world networks. In the case of biological networks they help in predicting the evolution and history of biomolecule interactions, provided we map properly real networks into the corresponding graph models. In this paper, we show that for biological graph models many of the existing parameter estimation techniques overlook the critical property of graph symmetry (also known formally as graph automorphisms), thus the estimated parameters give statistically insignificant results concerning the observed network. To demonstrate it and to develop accurate estimation procedures, we focus on the biologically inspired duplication-divergence model, and the up-to-date data of protein-protein interactions of seven species including human and yeast. Using exact recurrence relations of some prominent graph statistics, we devise a parameter estimation technique that provides the right order of symmetries and uses phylogenetically old proteins as the choice of seed graph nodes. We also find that our results are consistent with the ones obtained from maximum likelihood estimation (MLE). However, the MLE approach is significantly slower than our methods in practice.

## 1 Introduction

Many biological processes are regulated at the level of interactions between protein molecules. In recent decades, the development of experimental and bioinformatic methods allow researchers to obtain and make publicly available growing amount of data of various biological mechanisms involving different proteins. The whole network of these events that can be found and reconstructed using various techniques is customarily summarized as protein-protein interaction (PPI) networks. The PPI networks are often described as undirected graphs with nodes representing proteins and edges corresponding to protein-protein interactions.

The proliferation of biological data, in turn, encouraged a study of a series of theoretical models to develop a deeper understanding of the evolution and structural properties of the network representation. However, proper fitting of the biological data and development of statistical tests to check the validity of the models are critical to take advantage of the theoretical developments in graph models. The parameter estimation remains as a challenging problem in biological networks like PPIs, mainly because most of the classical estimation procedures were developed for static networks, and not tailored to specific properties of biological data and its underlying dynamics. In this paper, we propose a parameter estimation scheme for biological data with a new perspective of symmetries and recurrence relations, and point out many fallacies in the previous estimation procedures.

Since the inference techniques are closely coupled with the arrival process, we assume that networks evolve according to the following duplication-divergence stochastic graph model.

### Duplication-divergence model (DD-model)

There is a wide agreement [1], originally stemming from Ohno’s hypothesis on genome growth [2], that the main mechanism driving evolution of PPI networks is the duplication mechanism, in which new proteins appear as copies of some already existing proteins in the network. This is supplemented by a certain amount of divergence of random mutations that lead to some differences between patterns of interaction for the source protein and the duplicated protein.

There are many variations of duplication-divergence models in the literature, although they have not yet been studied or compared systematically. In this work, we use the model suggested by Pastor-Satorras et al. in [3], recommended in several surveys [4, 5] as a good possible theoretical match for PPI networks.

The model of Pastor-Satorras et al. constructs the graph in the following way. Given an undirected, simple, seed graph 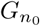 on *n*_0_ nodes and target number of nodes *n*, the graph *G*_*k*+1_ with *k* + 1 nodes^2^ evolves from the graph *G*_*k*_ a follows. First, a new vertex *v* is added to *G*_*k*_. Then the following steps are carried out:

- Duplication: Select an node *u* from *G*_*k*_ uniformly at random. The node *v* then makes connections to 𝒩_*k*_(*u*), the set of neighbors of *u* at time *k*.
- Divergence: Each of the newly made connections from *v* to 𝒩_*k*_(*u*) are deleted with probability 1 − *p*. Furthermore, for all the nodes in *G*_*k*_ except 𝒩_*k*_(*u*), create an edge from each of them to *v* independently with probability 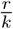.

The above process is repeated until the number of nodes in the graph reaches *n*. We denote the graph *G*_*n*_ generated from the DD-model with parameters *p* and *r*, starting from seed graph 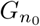, by *G*_*n*_ ∼ `DD-model` 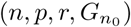 Note that the above model generalizes pure duplication model when *p* = 1, *r* = 0 [6, 7]. In some variations of the model (e.g. [8]), the nodes *v* and *u* will make a connection independently with a probability *q* that is much larger than *r/k*. However, the addition of *q* does not introduce significant changes in the properties of the graph, therefore we do not consider such a variation in this work.

### Motivation and contributions

In this work, we rigorously study the problem of parameter estimation in the duplication-divergence model using PPI datasets of seven species. The following points motivate this work and we present our key results with them.

### Symmetries of the graph

One important feature of the networks that was neglected in the previous studies is the distribution of the number of symmetries (formally known as automorphisms, which is the set of adjacency preserving permutations) generated by the fitted models with fine-tuned parameters. For the real-world PPI networks it turns out that the number of symmetries is considerably high, which is in stark contradiction with properties of many random graph models. For example, it is known that graphs generated from Erdős-Renyi model [9] and from preferential attachment model [10] are asymmetric with high probability. Therefore they cannot be reasonably justified as underlying generation schemes for PPI networks. We shall also see that the same phenomena may occur for the DD-model with some ranges of the parameters *p* and *r*.

Automorphisms are rarely studied in the context of biological networks and graph models. So far there are no theoretical results on automorphisms in the duplication-divergence model except the work in [11] for the limiting case of *r* = 0, where it was discovered that when both *p* = 0 and *p* = 1 the model produces graphs with a significant amount of symmetry.

Our main focus in this work is to take into account the number of automorphisms of the observed network to restrict the parameter search to a more meaningful range. Moreover, we show that most parameters outputted by previous estimation techniques fail to produce graphs having an order of automorphisms close to that of the PPI networks and therefore they are, in this regard, do not fit the DD-model well. We also note that cross-checking with the number of automorphisms of the real-world network forms a null hypothesis test for the model under consideration.

### Graph parameter recurrences

It is widely recognized that the asymptotics of structural properties such as the degree distribution and number of edges of the DD-model are crucial parameters, upon which judgment about the fitness of the model could be made. From the theoretical point of view, the analysis for the DD-model was presented in [12, 13], supporting the case that graphs derived from this model exhibit (under certain assumptions) power-law-like behavior. Moreover, the frequency of appearance of certain graph structures called graphlets (small subgraphs such as triangles, open triangles, etc.) can be viewed as another criterion for model fitting (see [4, 14] and the references therein). The triangles and *wedges* (paths of length 2 or star with two nodes) are particularly crucial as they are directly related to the network clustering coefficient. The high value of this coefficient is recognized in general as a significant characteristic of some biological networks including PPI networks [15], differentiating these networks from those which can be obtained, for example, from Erdős-Renyi or preferential attachment models.

Our approach is based on recreating graph evolution from a single snapshot of the observed network. We apply, for the first time, rigorous analyses to estimate parameters with the recurrence relations of degree and the number of wedges and triangles, recreating the dynamic process of DD-model construction. The advantages of this approach are twofold: first, the use of accurate iterative formulas allow us to achieve more realism and precision for finite graphs, which is in contrast to most of the previous studies that derive parameter estimates exclusively in terms of steady-state behavior. Second, it is not proven whether the steady state for such models even exists and whether the whole random process converges asymptotically (see Section 4 for more details).

### Maximum likelihood method

To substantiate the accuracy of our estimation technique, we apply the maximum likelihood method (MLE) with the importance sampling to the parameter inference problem, adapting the work of Wiuf et al. [16] to the DD-model. It turns out that the results of the MLE method are very similar to those that derived from our estimation method, and the estimated parameters in both the techniques generate data that is consistent with the observed network in terms of the number of automorphisms.

However, the MLE algorithm has much larger computational complexity, 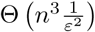 compared to 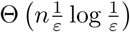 needed by our approach based on the recurrences (*n* and *ϵ* being the number of nodes and required resolution). Therefore, the analysis of graphs using MLE method is significantly slower for networks in practice and its application is impractical for networks exceeding 1000 nodes, which includes most of the real-world PPI networks. Furthermore, in the case of MLE, a large set of parameter values maximizes the likelihood function when the true *p* value is close to one, thus making it less reliable (see Section 6.1).

### Seed graph choice

It is well known that seed graphs play an important role in biological networks, and its improper selection will affect the estimated parameters of the fitted model [17, 5]. In particular, the task of determining the suitable range of parameters for the duplication-divergence model is always done under assumptions concerning the seed graph. In this work, we improve on the existing solutions by choosing the seed graph on the basis of phylogenetic ages of the proteins in the PPI data – the oldest proteins forms the seed graph. Although such a choice of seed graph is completely absent in prior literature, it is a natural pick as the seed graph itself is defined as the network that is comprised of the oldest entities in the given network. Our experiments in Section 6 show that seed graphs with oldest proteins provide better parameter estimates for the DD-model.

We made publicly available all the code and data of this project at https://github.com/krzysztof-turowski/duplication-divergence.

### Outline of the paper

In Section 2 we describe the PPI datasets that are considered in this paper. The influence of parameters of the DD-model on graph symmetries and the *p*-value calculation for comparing with the observed graph are given in Section 3. In Section 4 we provide a critique of various deficiencies of previous approaches to PPI networks parameter estimation, like lack of symmetries and overemphasis on power-law behavior. Section 5 describes our approach based on both automorphisms counting and exact iterative formulas for certain graph statistics. In this section we also present an MLE algorithm for parameter inference and compare methods in terms of their complexity and practical usage. Section 6 contains numerical results for both synthetic data generated from the DD-model and real-world PPI networks. Section 7 concludes the paper with a discussion of obtained results and their significance.

Table 1 provides the list of main notations used in the paper.

**Table 1:**
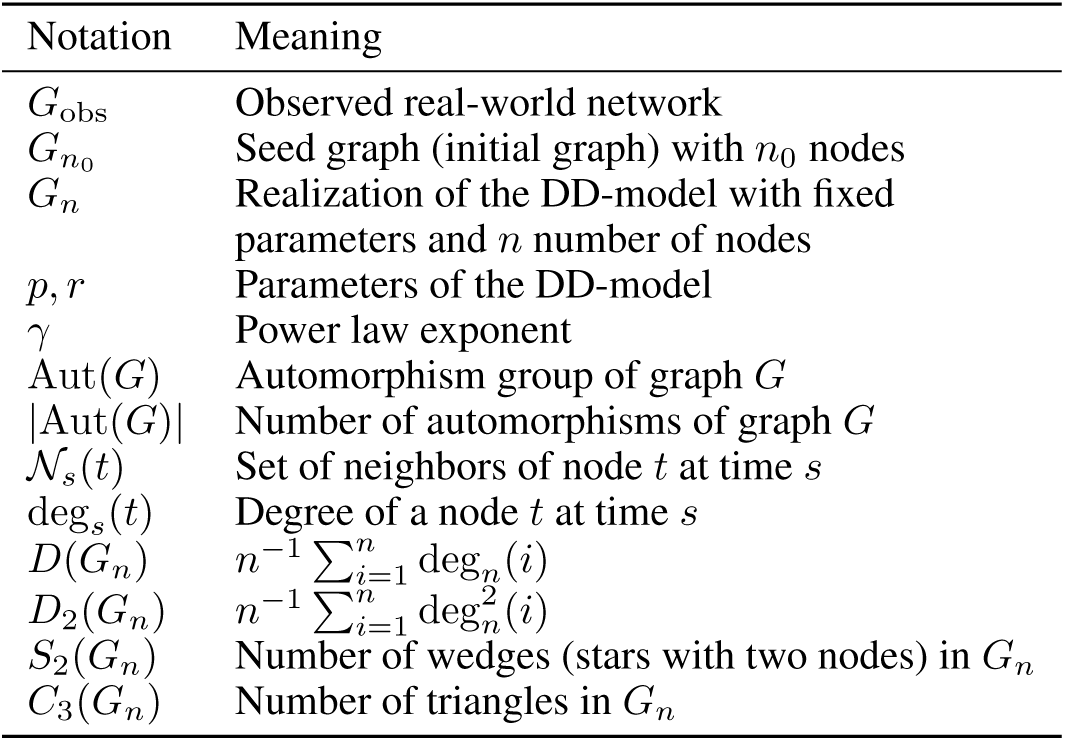
List of main notations

## 2 Datasets

We use protein-protein interaction networks (PPI) to verify the estimation techniques proposed in this paper. The data is collected from the BioGRID^3^, a popular curated biological database of protein-protein interactions. The networks formed from protein-protein interaction data are further cleaned by removing self-interactions (self-loops), multiple interactions (multiple edges), and interspecies (organisms) interactions of proteins. Thus the considered PPI networks only have physical and intra-species interactions. Unlike some of the previous studies that consider only the largest connected component, the DD-model we focus in this work incorporates disconnected subgraphs and isolated nodes.

Table 2 shows the different PPI datasets considered in this paper. We have also listed the logarithm of the number of automorphisms in the original graph, obtained using a publicly available program `nauty` [18]. We note here that the PPI dataset is growing as new interactions getting added on every new release of the dataset. Many previous studies were using older and less complete versions of the data, and therefore it is important to repeat the estimation procedures from those studies and compare them to our methods.

**Table 2:** Statistics of PPI networks used in this paper and the generated seed graph 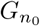 with nodes of the largest phylogenetic ages.

| Organism | Scientific name | Original graph $G_{\text{obs}}$ | | | Seed graph $G_{n_0}$ | |
| --- | --- | --- | --- | --- | --- | --- |
| | | # Nodes | # Edges | $\log \vert \text{Aut}(G)\vert $ | # Nodes | # Edges |
| Baker’s yeast | *Saccharomyces cerevisiae* | 6,152 | 531,400 | 267 | 548 | 5,194 |
| Human | *Homo sapiens* | 17,295 | 296,637 | 3026 | 546 | 2,822 |
| Fruitfly | *Drosophila melanogaster* | 9,205 | 60,355 | 1026 | 416 | 1,210 |
| Fission yeast | *Schizosaccharomyces pombe* | 4,177 | 58,084 | 675 | 412 | 226 |
| Mouse-ear cress | *Arabidopsis thaliana Columbia* | 9,388 | 34,885 | 6696 | 613 | 41 |
| Mouse | *Mus musculus* | 6,849 | 18,380 | 7827 | 305 | 7 |
| Worm | *Caenorhabditis elegans* | 3,869 | 7,815 | 3348 | 185 | 15 |

### 2.1 Selection of seed graph 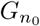

Previous studies typically assume the seed graph 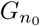 as the maximal clique (or the largest two cliques) in the graph *G*_*n*_ [5, 4]. Here we consider a novel formulation for the seed graph. We select the seed graph as the graph induced in the PPI networks by the oldest proteins. That is, the proteins in the observed PPI data that are known to have the largest phylogenetic age (taxon age). It is reasonable to expect that the same protein which appeared over different species also appears in their common ancestor. Hence proteins shared across many different, distant species are supposed to be older than others.

More precisely, the age of a protein is based on a family’s appearance on a species tree, and it is estimated via protein family databases and ancestral history reconstruction algorithms. We use Princeton Protein Orthology Database (PPOD) [19] along with OrthoMCL [20] and PANTHER [21] for the protein family database and asymmetric Wagner parsimony as the ancestral history reconstruction algorithm. These algorithms can be accessed via ProteinHistorian software [22].

Table 2 also lists the statistics of seed graphs 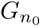 for different PPI networks. Even if the original PPI network is connected, the DD-model under consideration allows a disconnected graph to be the seed graph. Thus, similar to the formation of the PPI network, we consider the graph induced by oldest proteins including isolated nodes and disconnected subgraphs, not restricting ourselves to a connected component that introduces biases in the results.

## 3 Influence of parameters on symmetries of the model

For certain range of values of the parameters *p* and *r* of the DD-model, given *n* and 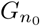, we show in this section that the model generates virtually only asymmetric graphs. However, we can put forward a question: are there any values of parameters that will yield graphs with the number of automorphisms is close to the real-world PPIs?

In Figure 1, we present the average number of symmetries in the logarithmic scale for graphs with different sizes. generated from the DD-model with a fixed set of parameters. As *p, r* → 0 or when *p* becomes very close to 1 we observe significantly larger values for the average number of automorphisms (since the generated graphs tend to have numerous isolated nodes or they become closer to a complete graph). For instance in Figure 1a, *p* = 1, *r* = 0.4 has 𝔼 [log Aut(*G*_*n*_)] = 1114, and *p* = 0, *r* = 0.4 has 𝔼 [log Aut(*G*_*n*_)] = 1253. But for large ranges of *p* and *r*, it is practically impossible to generate a graph exhibiting any noticeable symmetries. For example, *p* = 0.2, *r* = 2.4 has 𝔼 [log Aut(*G*_*n*_)] = 3.2; *p* = 0.6, *r* = 0 has 𝔼 [log Aut(*G*_*n*_)] = 1.3; and *p* = 0.4, *r* = 2.4 has 𝔼 [log Aut(*G*_*n*_)] = 0.12. These observations are consistent for different *n* and 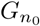 too, though the specific range of values of parameters will obviously change.

**Figure 1:**
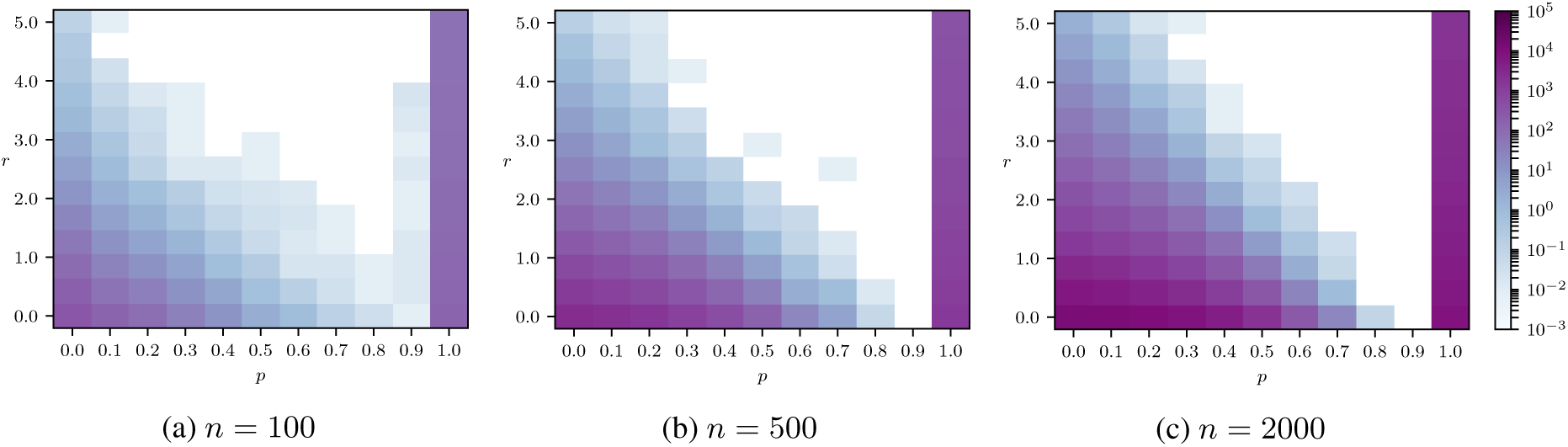
Logarithm of the expected number of automorphisms of graphs generated from the DD-model. The seed graph 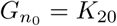.

The number of automorphisms in the DD-model behaves differently as compared to many other graph models including preferential attachment and E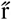dos-Rényi models. The preferential-attachment graphs are asymmetric (no nontrivial symmetries) with high probability when the number of edges a new node brings into the graph exceeds 2 [10], and almost every graph from the E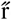dos-Rényi model is asymmetric [9]. On the other hand, the DD-model exhibits a large number of symmetries and it grows with the number of nodes, as shown in Figure 1.

These findings allow us to argue that only certain subsets of (*p, r*) pairs correspond to the expected number of automorphisms in the order of the required value. This means that it can be reasonably used as a falsification tool to discard certain parameter ranges and to verify parameter estimation methods. We provide a simple statistical test for checking the possibility of generating the required number of symmetries with the estimated parameters.

### Statistical test for significance of the number of symmetries with the estimated parameters

Given the real-world network *G*_obs_, seed graph 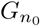, and the estimated parameters 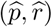 of the DD-model, we can estimate the statistical significance of the estimates with respect to the number of symmetries in *G*_obs_ as follows. Let 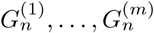 be *m* graphs generated from `DD-model` 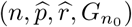. Then the *p*-value is now calculated as follows:

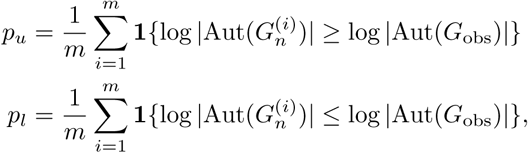

with **1** {*A*} as the indicator function of the event *A*. Then *p*-value = 2 min {*p*_*u*_, *p*_*l*_}. As an example, for a fixed parameter set, the empirical distribution of log |Aut(*G*)| is shown in Figure 2. The distribution is symmetrical and this justifies use of the symmetrical definition of *p*-value. A lower *p*-value indicates that the estimated parameters do not fit the observed network, and a higher value gives an argument for the estimated parameters being in agreement with the number of symmetries in *G*_obs_.

**Figure 2:**
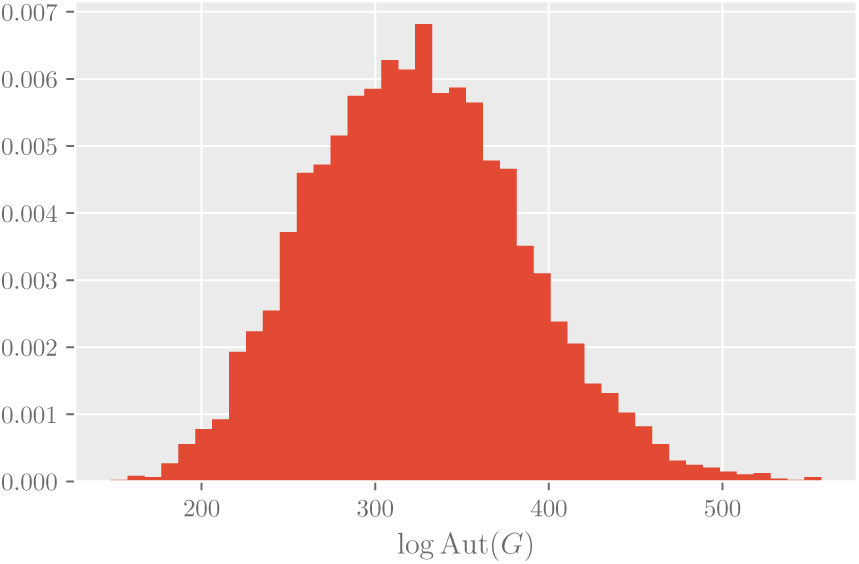
Normalized histogram of logarithm of number of automorphisms when *G*_*n*_ ∼ `DD-model`(500, 0.3, 0.4, *K*_20_).

## 4 Parameter Estimation and Why Existing Methods Fail in Practice?

Previous methods for the parameter estimation problem in the DD-model was first sketched in [3] and then considered more rigorously in [12]. Later, [5, 4] provided some extensions to the estimation procedures using the mean-field approximation of the average degree *D*(*G*_obs_) together with the steady-state expression of the power-law exponent *γ* of the degree distribution. Then, the values of *p* and *r* are computed, respectively, from the formulas:

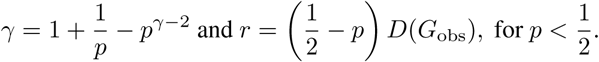

Table 3 presents the estimates of parameters *p* and *r* using the above method. Additionally, we present the average logarithm of the number of automorphisms computed from 10,000 graphs generated from the DD-model with the estimated parameters.

**Table 3:** Estimated parameters of the DD-model and average number of symmetries using mean-field approach.

| Organism | $\hat{p}$ | $\hat{r}$ | $\mathbb{E}[\log \vert \text{Aut}(G_n)\vert ]$ | $p$-value |
| --- | --- | --- | --- | --- |
| Baker's yeast | 0.28 | 38.25 | 0 | 0 |
| Human | 0.43 | 2.39 | 10.81 | 0 |
| Fruitfly | 0.44 | 0.75 | 3771.99 | 0 |
| Fission yeast | 0.46 | 1.02 | 897.48 | 0 |
| Mouse-ear cress | 0.44 | 0.43 | 18596.72 | 0 |
| Mouse | 0.48 | 0.12 | 34961.69 | 0 |
| Worm | 0.47 | 0.14 | 15700.26 | 0 |

### Mismatch in the number of symmetries and graph statistics

Comparing Tables 2 and 3, we observe that the number of symmetries of the graphs which are generated by the DD-model with parameters estimated via the mean-field approach differs significantly with that of the real-world PPI networks. Moreover, the estimated *p*-values are consistently zero for all the species because the observed values of the parameters under investigation fall far outside the range of the empirical distribution of the parameters for synthetic graphs generated with estimated *p* and *r*. This shows that the previously established estimation methods of the DD-model fail to capture the critical graph property of automorphisms, and thus do not fit the PPI networks accurately.

As shown in Table 2, the PPI networks exhibit some significant amount of symmetry, but far less than the maximum possible value (equal to *n* log *n*), which is attained when every node can be interchanged with every other node. This observation, along with the *p*-value test in Section 3, allow us to discard not only many models which produce only asymmetric graphs with high probability (such as Erdős-Renyi or preferential attachment model), but also effectively stands as a hypothesis test to verify that the fitting obtained by an estimation procedure can be safely assumed to match the model underlying real-world structures.

Similarly, for certain graph statistics *D*(*G*_*n*_), *S*_2_(*G*_*n*_) and *C*_3_(*G*_*n*_) (see Table 1 for notation), which are considered later in Section 5.1 for deriving our methods, we observe from Table 4 that the estimated parameters do not yield graphs that have the considered statistics close to the observed graph. Here the *p*-values are calculated in an equivalent way of number symmetries, just that now it is computed with respect to the graph statistics.

**Table 4:** Comparison of certain graph statistics of the observed graph and that of the synthetic data with parameters estimated via the mean-field approach.

| Organism | $D(G_{\text{obs}})$ | $\mathbb{E}[D(G_n)]$ | $p$-value | $S_2(G_{\text{obs}})$ | $\mathbb{E}[S_2(G_n)]$ | $p$-value | $C_3(G_{\text{obs}})$ | $\mathbb{E}[C_3(G_n)]$ | $p$-value |
| --- | --- | --- | --- | --- | --- | --- | --- | --- | --- |
| Baker's yeast | 172.76 | 115.10 | 0 | 220.35M | 45.33M | 0 | 9.77M | 370.49K | 0 |
| Human | 34.30 | 19.39 | 0 | 52.25M | 7.02M | 0 | 1.07M | 105K | 0 |
| Fruitfly | 13.11 | 7.87 | 0 | 2.94M | 1.45M | 0 | 195.96K | 77.61K | 0 |
| Fission yeast | 27.64 | 6.72 | 0 | 7.42M | 215.84K | 0 | 223.61K | 1.14K | 0 |
| Mouse-ear cress | 7.39 | 2.23 | 0 | 2.98M | 44.46K | 0 | 23.34K | 23.27 | 0 |
| Mouse | 5.35 | 0.82 | 0 | 2.95M | 9.33K | 0 | 10.22K | 0.79 | 0 |
| Worm | 4.04 | 0.90 | 0 | 346.13K | 5.32K | 0 | 2.41K | 0.49 | 0 |

We would like to point out several other deficiencies in the known estimation procedures, which could be the reasons behind such a divergence between the number of symmetries of the PPI networks and its proposed theoretical model.

### Power-law behavior

The parameter estimation of the DD-model introduced in prior works, such as the one that was presented at the beginning of this section, assumes that the PPI networks are scale-free. This property, stating that the degree distribution of the PPI networks is heavy-tailed or, more precisely, that the number of vertices of degree *k* is proportional to *k*^−*γ*^ for some constant *γ* > 0 [23, 24]. With this assumption, some (see [8] for example) argue that the estimated value of the exponent for the PPI networks satisfies 2 *< γ <* 3. However, there are some serious counterarguments to this claim, and it is challenged on statistical grounds that the PPI graphs do not fall into the power-law degree distribution category [25, 26].

To each of the PPI networks in Table 2, we fit the coefficients of power-law distribution with the cut-off following the methodology of Clauset et al. [27]. We note here that cutoff is required in all the cases since the power-law behavior mostly happens in the tails of the degree distribution. However, we find that the cutoff neglects a huge percentage of the data. For example, for a fitting of baker’s yeast PPI network, as shown in Figure 3, the cutoff is 582, which is at 94.98 percentile of the degree data, i.e., the power-law fitting does not take into account 94.98% of the data. With the cutoff and the percentiles of all the species listed in Table 5, we remark that any method to estimate the parameters *p* and *r* involving power-law exponent need not result in good approximations since it discards data from a majority of the network.

**Table 5:** Estimated power law exponent and required cutoff percentile with the mean-field approach

| Organism | $\hat{\gamma}$ | Cutoff percentile |
| --- | --- | --- |
| Baker's yeast | 4.55 | 94.98 |
| Human | 2.85 | 92.33 |
| Fruitfly | 2.71 | 88.00 |
| Fission yeast | 2.43 | 88.31 |
| Mouse-ear cress | 2.68 | 93.89 |
| Mouse | 2.29 | 78.58 |
| Worm | 2.41 | 88.23 |

**Figure 3:**
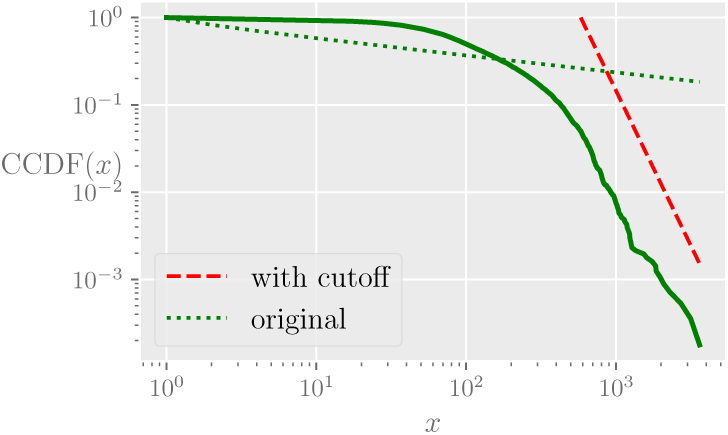
Complementary cumulative distribution function (CCDF) of baker’s yeast and power law fitting.

### Steady state assumption

Previous research on the DD-model, both on the level of theoretical analysis of the model properties and the level of parameter estimation of real-world PPI networks, focus heavily on the asymptotic and steady-state behavior [3]. Most of the previous results on the functional form of certain graph statistics in the DD-model are under the strong assumption of steady-state [28, 8]. But they do not provide any theoretical proof for convergence to steady-state. Moreover, these steady-state asymptotic results, even when achievable, do not give any bounds on the rate of convergence. This, in turn, raises questions about the straightforward applicability of such theoretical results to parameter estimation.

The previously used methods of parameter estimation also suffer from another issue: for simplicity, they assume that the average degree of the network does not change during the whole evolution from 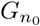 to *G*_*n*_. This is not only highly implausible in practice, but also impose direct relation between *p* and *r*, and hides any dependency that might be discovered from various properties of the networks.

### Seed graph choice

As shown by previous studies (most notably in Hormozidari et al. [5]), choice of the seed graph plays a significant role in graph evolution, directly contributing to the order of growth of many important graph statistics.

The seed graph is typically assumed to be the largest clique (or a connected graph of the largest two cliques) of the observed graph. Then random vertices and edges are gradually added to the network, preserving the average degree of the final network, to make the size of the network to a fixed value of *n*_0_. This method is motivated by the infinitesimally small probability with which there could appear a clique of a greater size during graph evolution. Such a procedure has no formal theoretical guarantees and does not have any clear justification from a biological perspective [5].

Our natural approach to select the seed graph is based on the extra-network information about the estimated age of proteins, described in Section 2.1.

## 5 Main results

Our main constructive results concern the relation between the parameters of the model and the number of symmetries exhibited by graphs generated from it. Additionally, we present two parameter estimation algorithms, one based on recurrences characteristic for certain graph statistics, the other based on the well-known maximum likelihood approach.

### 5.1 Our method: parameter estimation using recurrence relations

Our basic tool to infer the parameters of DD-model for a given the PPI network is a set of the exact recurrence relations for basic graph statistics, which relate their values at time *k* and *k* + 1 of graph evolution. Such recurrence relations are sufficient to estimate model parameters, as the whole sequence of graphs from the initial graph 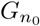 to the final graph *G*_*n*_ can be split into steps consisting of the addition of a single vertex and the changes introduced by the added vertex.

Typically, five statistics of the random graph *G*_*n*_ are studied in literature: number of edges *E*(*G*_*n*_), mean degree of the network 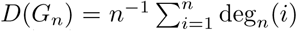, mean squared degree 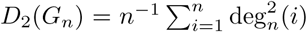, number of triangles (3-cliques) *C*_3_(*G*_*n*_), and number of *wedges S*_2_(*G*_*n*_) (wedges are also called 2-stars or paths of length 2 in prior literature, and number of wedges includes counts of triangles and open triangles).

However, for every graph *H* on *n* vertices it is true that 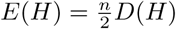 and 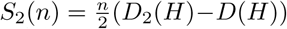. Therefore, it is sufficient to analyze only the three of above-mentioned graph statistics: *D*(*n*), *S*_2_(*n*) and *C*_3_(*n*).

As a first step, we derive the following recurrence relations for the chosen statistics.

#### Theorem 1.

*If G*_*n*+1_ *∼ `DD-model`*(*n* + 1, *p, r, G*_*n*_), *then*

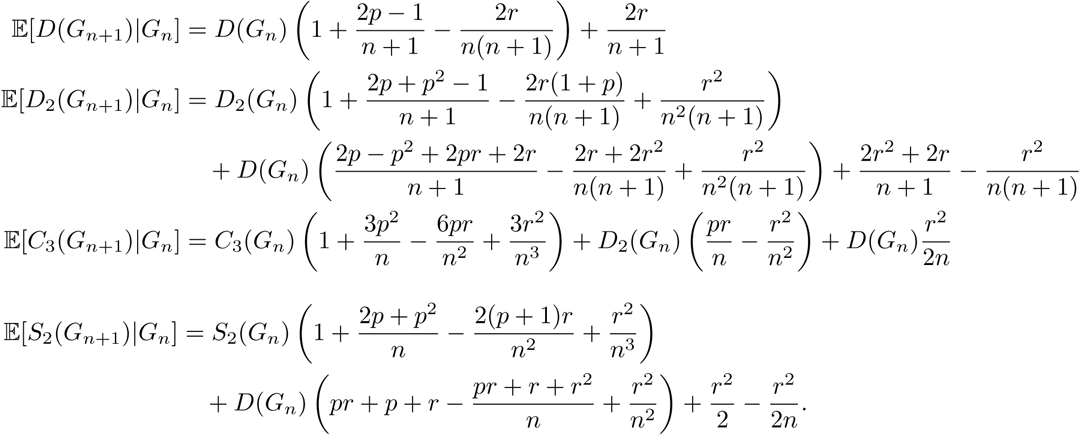

*Proof*. See Appendix B

In Figure 4, we verify Theorem 1 by comparing 𝔼 [*D*_*n*_], for various *n*, computed using theory and experiments.

**Figure 4:**
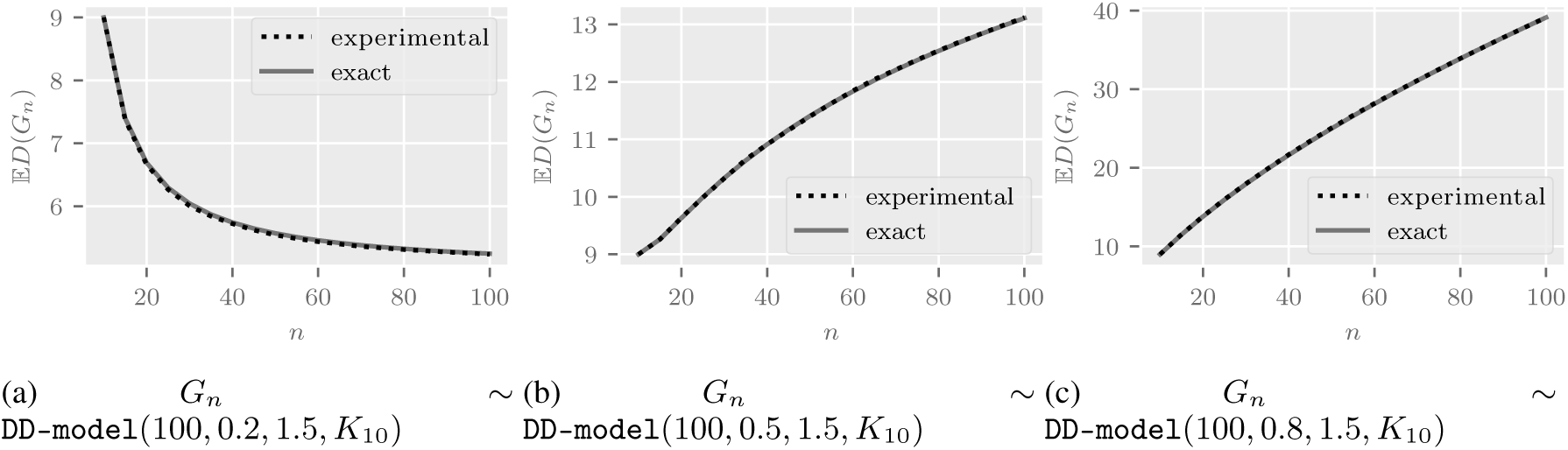
Comparison of 𝔼 [*D*(*G*_*n*_)] vs *n* computed via Theorem 1 and via experiments.

The expressions given in Theorem 1 implicitly define a function 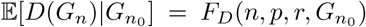, which is a cornerstone of our algorithm. Similar functions exist for recurrences based on other statistics of 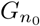 and *G*_*n*_. Now we claim that the result of Theorem 1 in terms of expectation can be used for the graph statistics with high probability too. Figure 5 shows the concentration of empirical distribution of different graph statistics.

**Figure 5:**
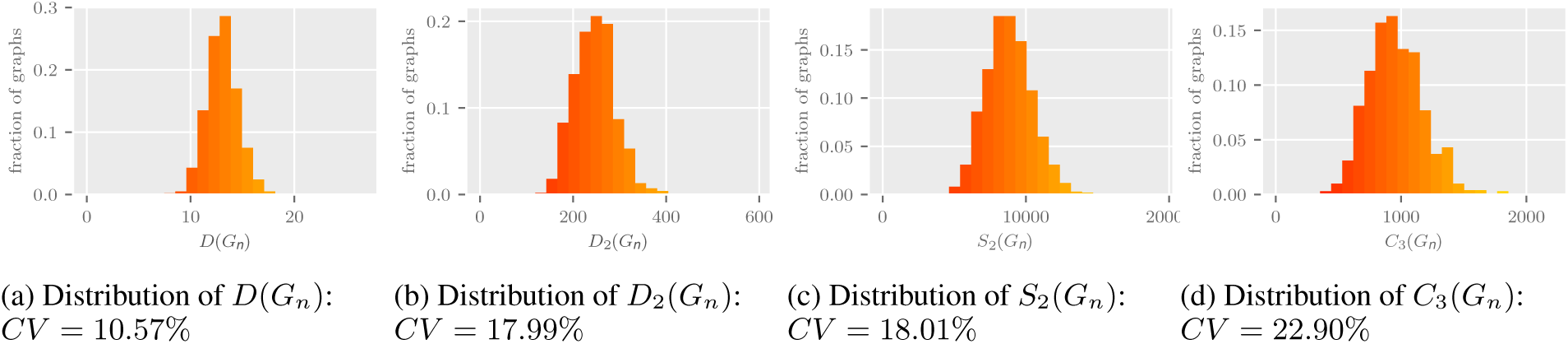
Empirical distribution of graph statistics: *G*_*n*_ ∼ `DD-model`(100, 0.5, 1.5, *K*_10_). Coefficient of variation *CV* is defined as the ratio of empirical standard deviation and empirical mean. The lower values of *CV* in the sub-figures show the concentration of the considered graph statistics.

Although we don’t need an explicit formula for *F*_*D*_ in our algorithm, we may derive one from the recurrences:

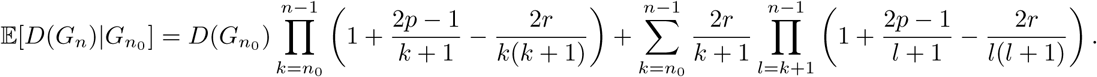

Although this is outside of the scope of this article, we note that such an expression allows us to find, for example, the asymptotic order of growth for *D*(*G*_*n*_) and for other statistics.

Though closed form solution of recurrences with *G*_*n*_ and 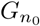 could be difficult to obtain, Theorem 1 is sufficient to formulate an efficient algorithm for finding the parameters of the model. The crucial feature is that all parameters are monotonic, that is, the larger the parameters *p* and *r*, the larger the values of *D*(*G*_*n*_) and other statistics.

Algorithm 1 presents our estimation technique for finding 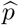 with the recurrence relation for *D*(*G*_*n*_) (which will be *D*(*G*_obs_) when we consider real-world network), assuming 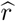 is known beforehand. However, sufficient number of samples of 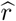 from the interval [0, *n*_0_] is adequate to get a feasible solution set of 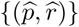 with a desired resolution. The algorithm also works for recurrence relations of *S*_2_(*G*_*n*_) and *C*_3_(*G*_*n*_) with evident modifications.

#### Algorithm 1 Parameter estimation via recurrence relations of *D*(*G*_*n*_).

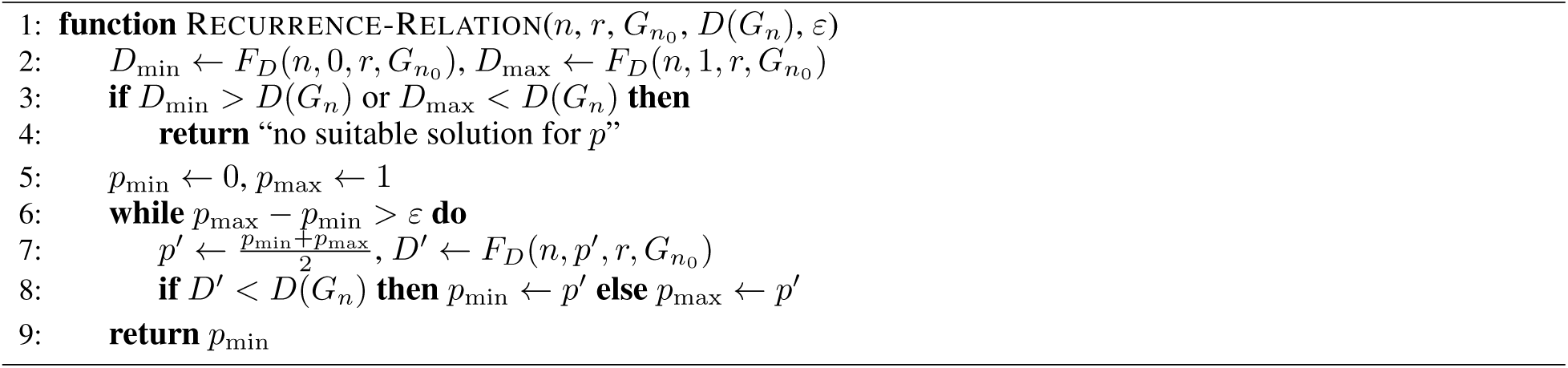

We note here that for each graph property under consideration, *D, S*_2_ or *C*_3_, the estimation algorithm returns a curve (more precisely, a set of feasible points). Now, if we find a concurrence in the solutions to the recurrence relations of various graph statistics, we know that a necessary condition for the presence of duplication-divergence model has been satisfied. On the other hand, if the curves were not having a common crossing point, it suggests that the DD-model may not be the appropriate fit for the observed network. We denote the above estimation procedure using the relations of all three graph statistics as the RECURRENCE-RELATION method.

### 5.2 Parameter estimation via maximum likelihood method

An alternative way of estimating parameters of the DD-model is the maximum likelihood estimation (MLE). With MLE, the estimated parameters 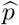 and 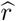 are given by max_*θ*=(*p,r*)_ *L*(*θ, G_n_*), where the likelihood function is *L*(*θ, G*_*n*_) is the probability of generating *G*_*n*_ from 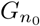 for fixed parameters *θ*, i.e.

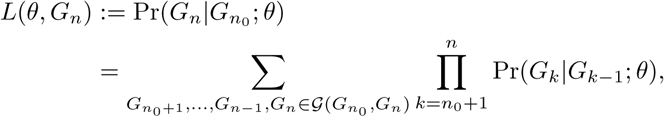

where 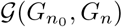 is the set of all sequences of graphs that starts with 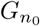 and ends at *G*_*n*_. Given a fixed sequence of graph evolution history 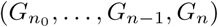, it is straightforward to calculate the likelihood, but *L*(*θ, G*_*n*_) requires summation over all histories, which has exponential number of possibilities. In [16], the authors present an importance sampling strategy to approximate the likelihood and thereby estimate the parameters. It is based on the idea of traversing backwards in history (*G*_*n*_ to *G*_*n*−1_ and *G*_*n*−1_ to *G*_*n*−2_ likewise) on one sample path of graph evolution sequence via Markov chain. We adapt their algorithm to our DD-model and the complete algorithm is presented in Supplementary Material.

We now provide a brief description of the importance sampling procedure. The idea is to express likelihood in terms of a known reference parameter *θ*_0_ instead of unknown *θ*. Now, the likelihood can be rewritten as an expectation with respect to *θ*_0_ and can be estimated via Monte Carlo simulations (see [16] for more details):

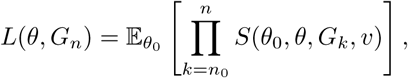

where

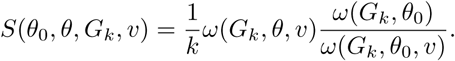

Here *ω*(*G*_*k*_, *θ, v*) is the probability of creating the graph *G*_*k*_ from *G*_*k*−1_ through the addition of a node *v*, with parameter as *θ. v* can be chosen as any node in *G*_*k*_ such that its removal would result in a positive probability *G*_*k*−1_ under the DD-model. The variable *ω*(*G*_*k*_, *θ*) is the transition probability *ω*(*G*_*k*_, *θ, v*), summed over all possible *v*. In fact, *ω*(*G*_*k*_, *θ, v*) itself is the normalized sum of *ω*(*G*_*k*_, *θ, v, w*), over all possible nodes *w*, which is the probability of producing a graph *G*_*k*_ from *G*_*k*−1_ by adding a node *v* that is duplicated from node *w*.

### 5.3 Computational complexity of parameter estimation methods

Let us now assume that the *n*_0_ is fixed and we are interested in results up to a resolution *ε*, that is, the values of *p* and *r* are stored in such a way that two numbers within a distance less than *ε* are indistinguishable.

For our algorithm 1, RECURRENCE-RELATION, a single pass requires 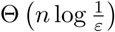, as it uses a binary search for *p* and for every intermediate value of *p* it executes exactly *n* − *n*_0_ steps of *for* loop, each requiring constant time. Now it is sufficient to sample 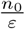 different values of *r*, therefore the total running time to find suitable 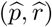 pairs is 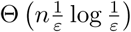.

On the other hand, the MLE algorithm needs to compute at every step values of the *ω* function for all possible pairs of *v* and *w* for each graph *G*_*k*_. This is the case because in DD-model every vertex *v* could be a duplicate of every other vertex *w* always with some non-zero probability at every stage of the algorithm. This means that we require Θ(*k*^2^) steps at each iteration of the algorithm; therefore 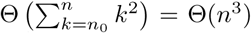 steps in total. Unfortunately, even clever bookkeeping and amortization is not much of a help here.

Additionally, we need to estimate the likelihood for each pair (*p, r*) independently, as maximum likelihood function does not have the monotonicity property, so it requires in total 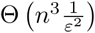 steps to find all feasible pairs up to a desired resolution of *ϵ*.

Moreover, as it was suggested by Wiuf et al. in [16], importance sampling provides good quality results only for at least 1000 independent trials. This adds up a constant factor not visible in the big-θ notation, but significant in practice, effectively making the algorithm infeasible for the real-world data without using supercomputer power.

## 6 Numerical results

In this section, we evaluate our methods on synthetic graphs and real-world PPI networks.

### Estimation of tolerance interval

We find the tolerance interval of the estimated *p* and *r* values for the fitted DD-model as follows. For a given network *G*_obs_ and a seed graph 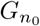, first the RECURRENCE-RELATION algorithm outputs a set of solutions 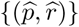. For each of the feasible pairs, we then estimate the confidence interval of the graph property with which RECURRENCE was calculated. For instance, if the property used was the empirical mean *D*, graph samples generated from `DD-model` 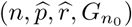 are used to estimate expectation 𝔼 [*D*(*G*_*n*_)] and variance Var[*D*(*G*_*n*_))] of *D*(*n*). A 95% confidence interval of *D*(*G*_*n*_) is then given by

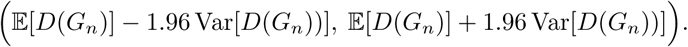

The Gaussian distribution assumption used in the above expression is indeed a good approximation for the distribution of *D* for large *n*. Now by fixing 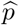, we can calculate a tolerance interval 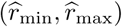 for the estimated parameter 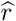. In the following experiments, for demonstrating the above approach, we focus on two graph statistics *D* and *C*_3_ (parameter estimation will include *S*_2_ too).

Our estimation procedure can be summarized follows:

- We employ the RECURRENCE-RELATION algorithm for solving graph RECURRENCEs of the three graph statistics *D, S*_2_ and *C*_3_, and we identify a set of solutions for *p* and *r*.
- With *G*_*n*_ ∼ `DD-model` 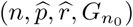, we find the tolerance interval of 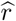 using the confidence interval of *D*(*G*_*n*_) and *C*_3_(*G*_*n*_).
- We look for crossing points of the plots in the figure, and the range of values of *p* and *r* where the confidence intervals meet around the crossing point. We call such a range of values as *feasible-box*.
- Though any point in the feasible-box is a good estimate of *p* and *r*, to improve the accuracy, we uniformly sample a fixed number of points from the box and choose the pair that gives maximum *p*-value with respect to the number of automorphisms of the given graph *G*_obs_.

### 6.1 Synthetic graphs

In this section, we derive insights by studying our method and MLE empirically on synthetic data. We generate two random graph samples 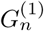 and 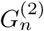 from the DD-model with the following parameters:

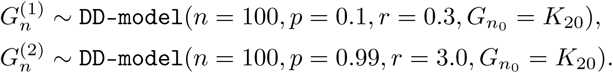

The choice of parameters in 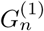 and 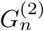 show different regimes in the following studies. Moreover the parameters are chosen in such a way that the generated graphs have non-trivial symmetries.

Figures 6a and 6b plot the sets of feasible points identified by the recurrence relations using RECURRENCE-RELATION method. The light shaded bands show the tolerance intervals of *r*. We observe that the crossing points and the tolerance intervals are fairly close to the original parameters. Figures 6c and 6d display the heat-plot of log-likelihood function of the MLE for different values of the parameters. The log-likelihood function maximizes at (*p, r*) pairs close to the original parameters, but not up to the resolution of RECURRENCE-RELATION method.

**Figure 6:**
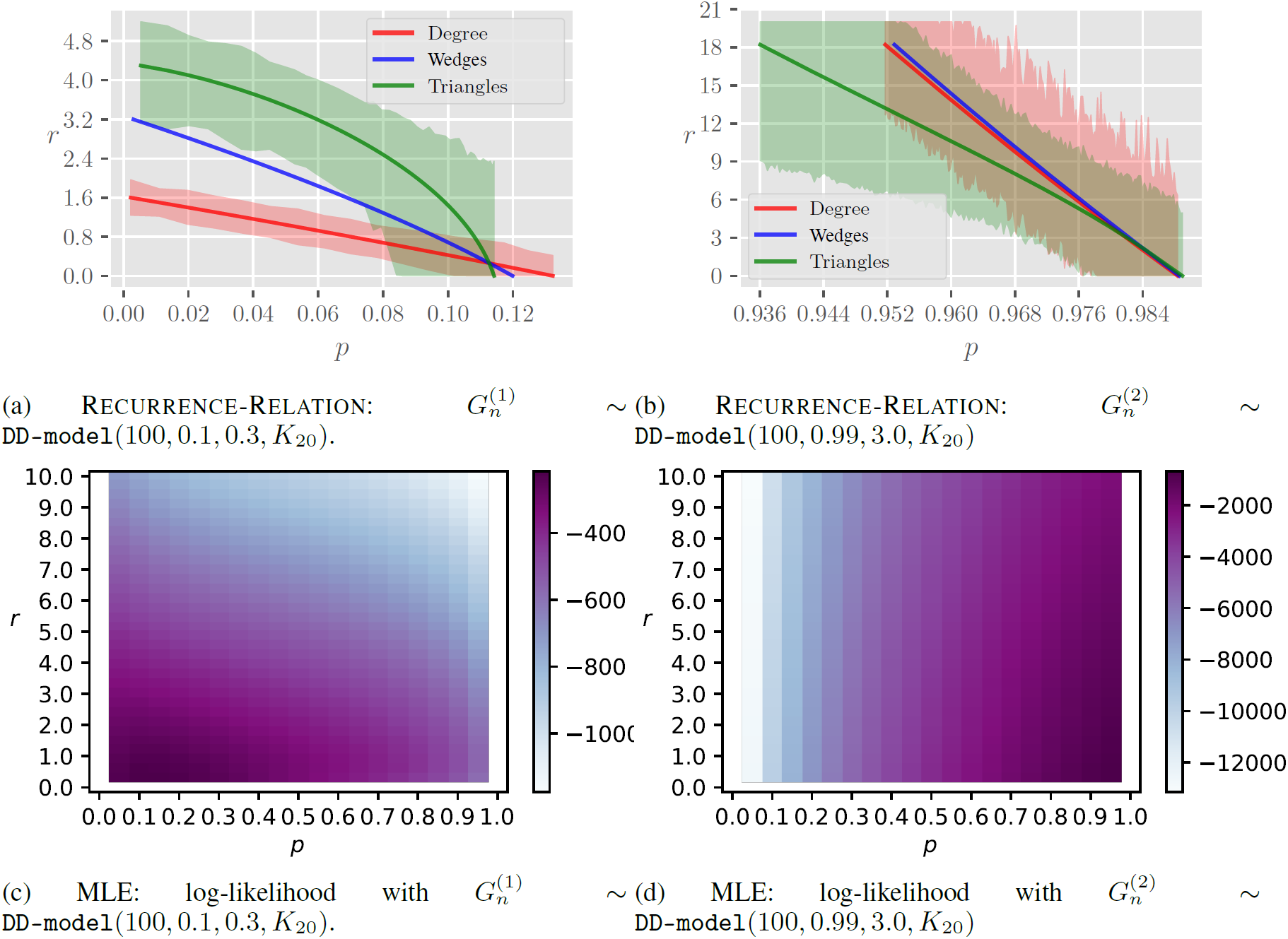
Results on synthetic networks: RECURRENCE-RELATION and maximum likelihood estimation (MLE) methods

In Table 6 we produce the statistical significance of the best estimated parameter pairs via both the RECURRENCE-RELATION and the MLE. The best pair is found in the RECURRENCE-RELATION method from 1000 uniform samples in the feasible-box centered at the point where the three curves are in agreement, and for the MLE, it is found from 1000 uniform samples in the maximum log-likelihood area if no unique maximizer exists. The estimates from both the techniques demonstrate the presence of the DD-model in the given graphs 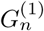 and 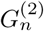 (*p*-value *>* 0.1), the best pair of RECURRENCE-RELATION estimator has much higher *p*-value and certainly outperforms MLE.

**Table 6:**
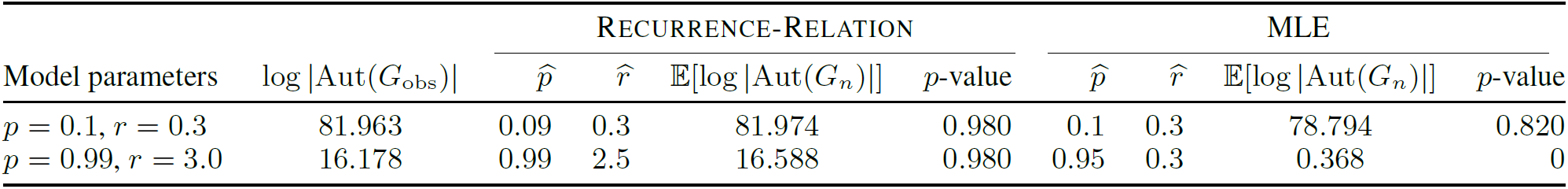
Results on synthetic networks: average number of automorphisms and *p*-value

We note that for the first graph 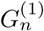 the results obtained by both methods are almost identical, in terms of 𝔼 [log |Aut(*G*_*n*_)|] and *p*-values. For the second graph 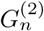, *the log-likelihood function of MLE is nearly flat for large values of p, and thus MLE returns less reliable estimates*. This in turn results in a larger deviation of the number of automorphisms from the observed graph. Our algorithm on the other hand provides a better estimate even when *p* is close to 1. To sum up, we find that our algorithm does not perform worse than MLE in terms of quality and achieves better performance than MLE when *p* is high. It also has much lower computational complexity.

### 6.2 Real-world PPIs

We apply recurrence-based estimator to PPI networks of seven species listed in Table 2. As mentioned in Section 2.1, the seed graph 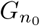 is assumed as the graph induced by the nodes having the largest phylogenetic age.

Figure 7 presents plots of RECURRENCE-RELATION estimator for seven species. In all the figures, the plots meet or come very close at a specific point. This illustrates the presence of the DD-model in all the considered species. Furthermore, Table 7 calculates the statistical significance of the fitted DD-model with respect to the number of automorphisms in the observed PPI networks. The estimated *p*-values are remarkably high and most often much larger than 0.4 (except in one case), demonstrating that the fitted DD-models exhibit symmetries closer to the real-world PPIs.

**Table 7:** Parameters of the real-world PPI networks estimated using RECURRENCE-RELATION method

| Organism | $\hat{p}$ | $\hat{r}$ | $\mathbb{E}[\log \vert \text{Aut}(G_n)\vert ]$ | $p$-value |
| --- | --- | --- | --- | --- |
| Baker's yeast | 0.98 | 0.35 | 293.27 | 0.71 |
| Human | 0.64 | 0.49 | 2998.81 | 0.51 |
| Fruitfly | 0.53 | 0.92 | 1073.83 | 0.64 |
| Fission yeast | 0.983 | 0.85 | 705.278 | 0.74 |
| Mouse-ear cress | 0.98 | 0.49 | 6210.36 | 0.13 |
| Mouse | 0.96 | 0.32 | 8067.56 | 0.67 |
| Worm | 0.85 | 0.35 | 3352.91 | 0.48 |

**Figure 7:**
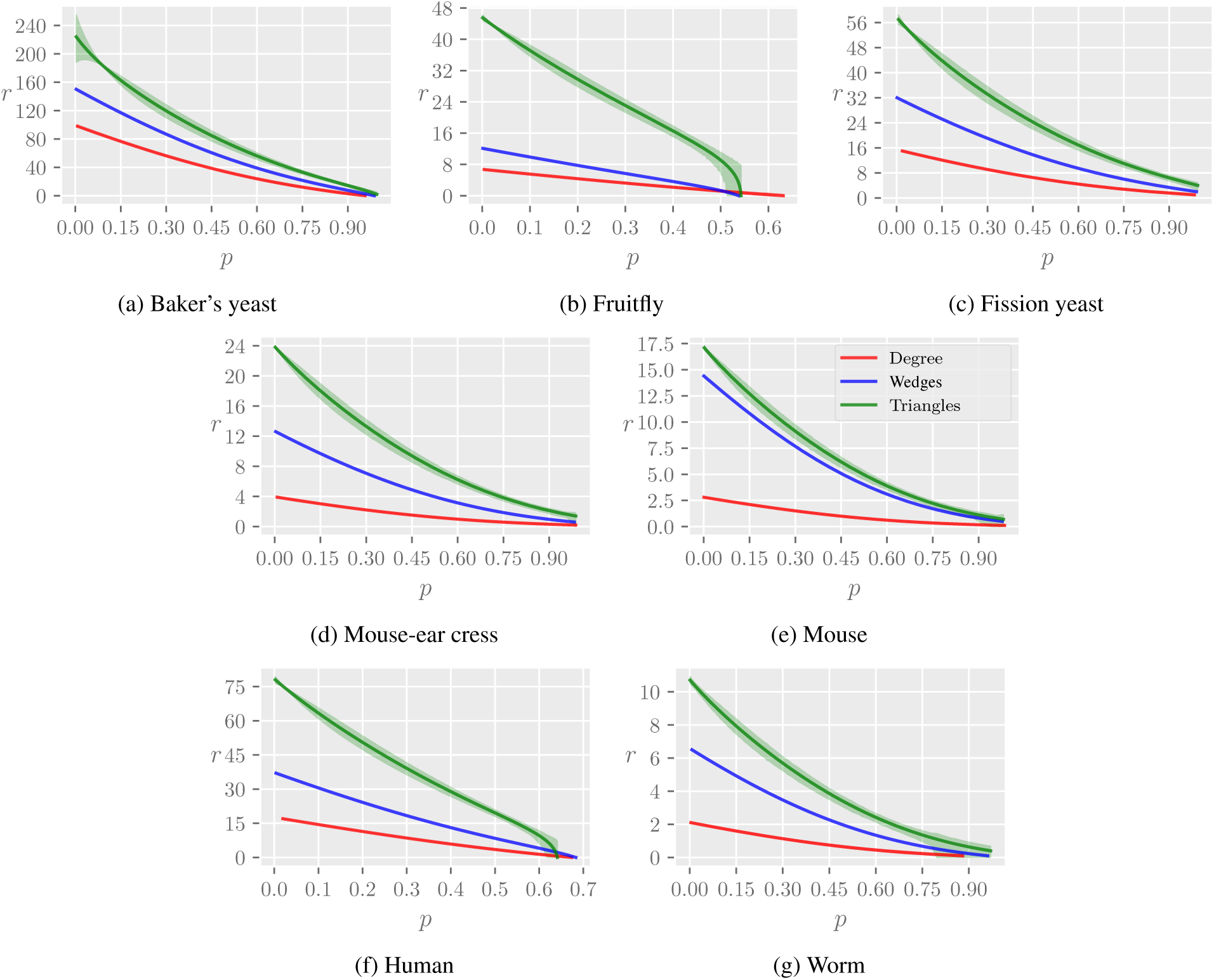
Results on PPI networks: RECURRENCE-RELATION method

## 7 Discussion and implications

We focus in this work on fitting dynamic biological networks to a probabilistic graph model, from a single snapshot of the networks. Our attention here is on a key characteristic of the networks – the number of automorphisms – that is often neglected in modeling. Using the number of automorphisms as a measure to sample parameters from the parameter space may raise serious questions about its practicality (like some slower maximum likelihood estimation methods for graph fitting). To address this, our approach in this paper to combine the number of symmetries with a faster method of recurrence relations, which allows us to narrow down the parameter search, finds high relevance in practice.

We argue that many existing parameter estimation techniques fail to take into account the number of symmetries of real-world networks, leading to serious concerns in the fitting methodology. Previous studies made unrealistic assumptions like the steady-state behavior of the model, and it could be the reason behind erroneous estimates. Our proposed fitting method based on exact recurrence relations with minimal assumptions works well on synthetic data and real-world protein-protein interaction (PPI) networks of seven species. We also formulate a simple statistical test in terms of the number of symmetries. Since the PPI networks are expanding with new protein-protein interactions getting discovered, we make sure to use up-to-date data so that the fitted parameters in this paper can serve as a benchmark for future studies.

We note here that the method introduced in this work is applicable to a variety of dynamic network models, as for many models there exist recurrence relations similar to the ones presented here. A systematic way of parameter estimation can also be seen as an introductory work to other important problems in biological networks. One example of such a problem is the *temporal order problem* [29]: given a network, the task is to recover the chronology of the node arrivals in the network. Parameter estimation provides us with better knowledge about the specific characteristics of the model that retains temporal information in its structure.

## 8 Acknowledgments

This work was supported by NSF Center for Science of Information (CSoI) Grant, and in addition by NSF Grants CCF-1524312, and NIH Grant 1U01CA198941-01.

## Appendix

### A Maximum likelihood algorithm

Function MAXIMUMLIKELIHOODVALUE, as shown below in Algorithm 2, presents a single pass of the MLE technique. Here *θ*_0_ = (*p*_0_, *r*_0_) is the initial parameter set which can be chosen with some extra knowledge of the given network or even arbitrarily; but with a proper choice of *θ*_0_, a faster convergence to the true value of likelihood is guaranteed.

We note here that the MLE procedure has to be run multiple times, and the average of the *L*’s (see algorithm) from the multiple runs gives an estimate of the likelihood at the inputted (*p, r*). The procedure needs to be repeated for all the relevant (*p, r*) pairs. The estimated parameters are the points for which the likelihood function is maximized. See Section 5.2 for details.

#### Algorithm 2 Single run for the likelihood value computation.

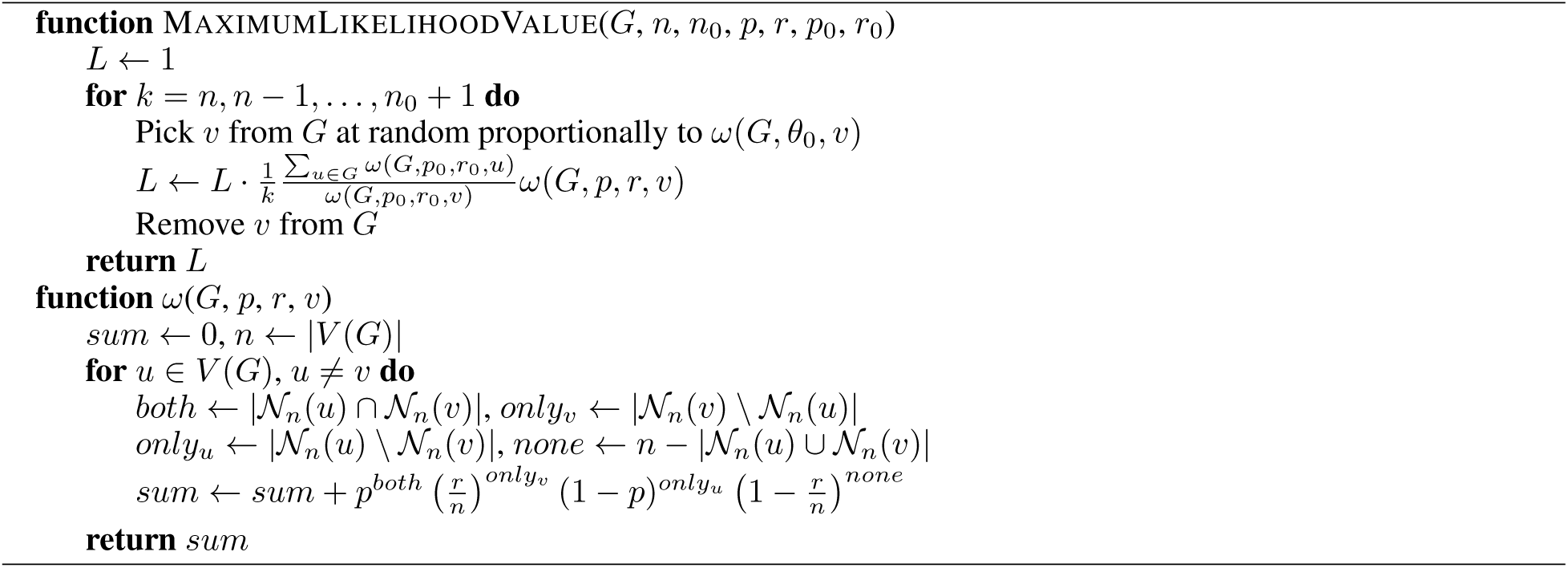

### B. Proof of Theorem 1

We recall Theorem 1.

**Theorem**. *If G*_*n*+1_ *∼ `DD-model`*(*n* + 1, *p, r, G*_*n*_), *then*

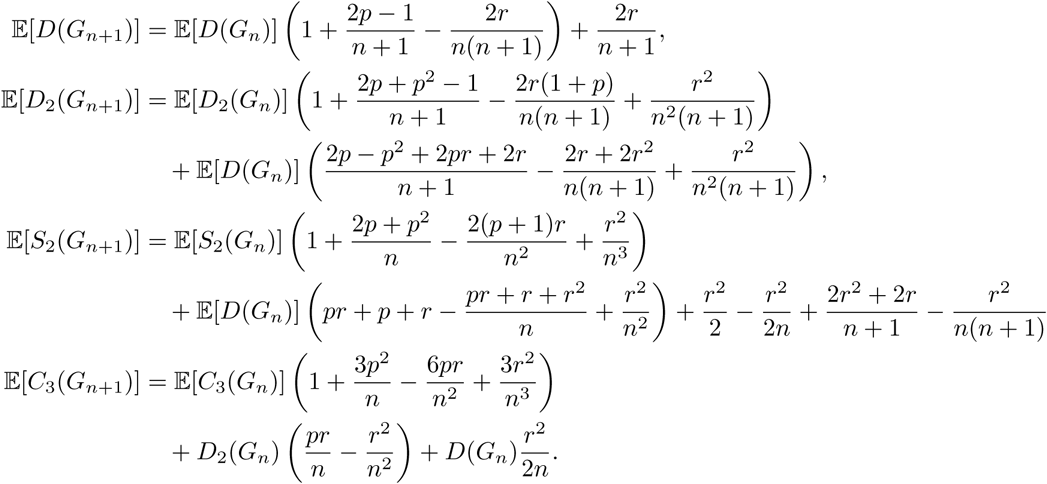

*Proof*. Let deg*t*(*s*) be the degree at time *t* of a vertex added at time *s*, which is same as node label *s*, and parent(*t*) be a vertex which was chosen from *G*_*t*−1_ for the duplication step at time *t*.

It follows from the definition of the model that degree of the new vertex *n*+1 is the total number of edges from the vertex *n*+1 to 𝒩_*n*_(parent(*n* + 1)) (each of which is formed from choosing nodes independently from 𝒩_*n*_(parent(*n* + 1)) with probability *p*) and to all other vertices (each of which is formed from nodes chosen independently from a set *V* (*G*_*n*_)\ 𝒩_*n*_(parent(*n* + 1)) with probability 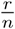).

It can be then expressed as a sum of two independent binomial variables:

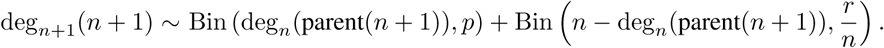

#### 1. Recurrence for D(*G*_*n*_)

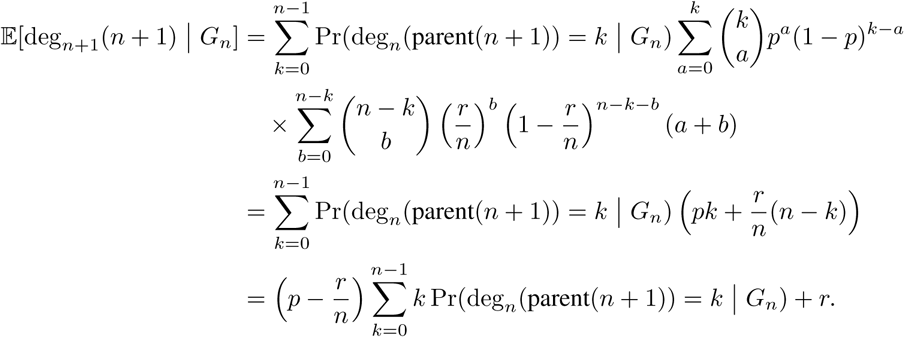

Since in the definition of the model it is stated that the parent is selected uniformly at random, we know that Pr(parent 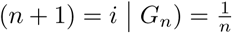 and therefore

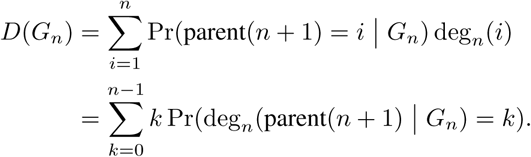

Combining the last two equations, we get

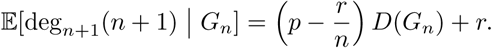

Using the above, we find the following recurrence for the mean degree of *G*_*n*+1_:

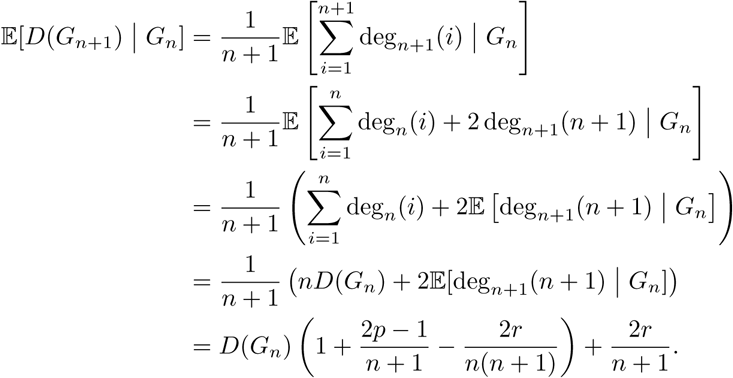

Now from the law of total expectation:

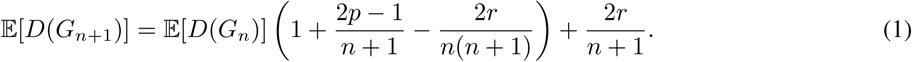

#### 2. Recurrence for *D*_2_(*G*_*n*_)

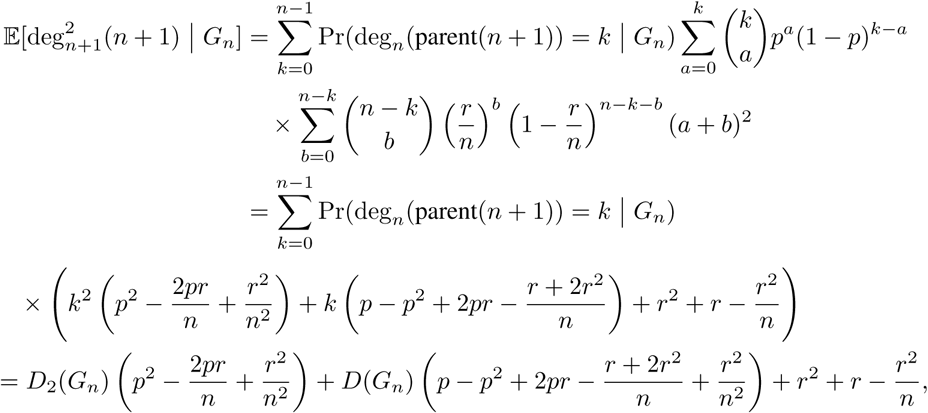

since we have, as before,

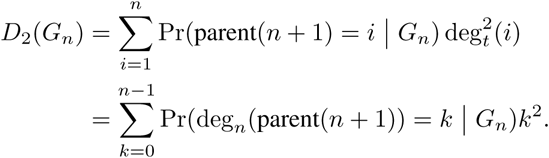

Now we proceed with the second moment of degree distribution of *G*_*n*_. Let *I*_*n*+1_(*i*) be an indicator variable whether there is an edge between *n* + 1 and *i*. Then the following basic results follows:

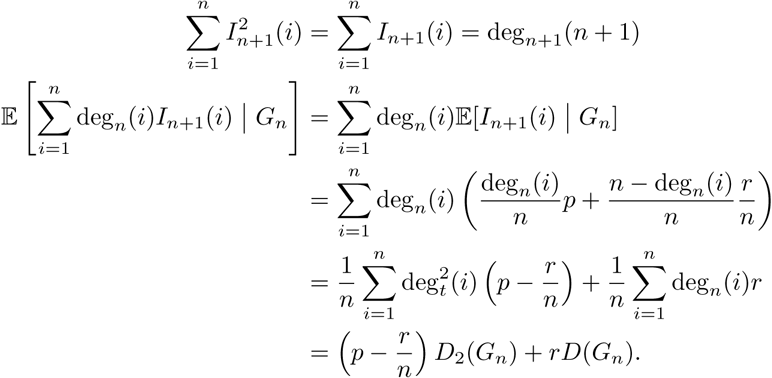

Now,

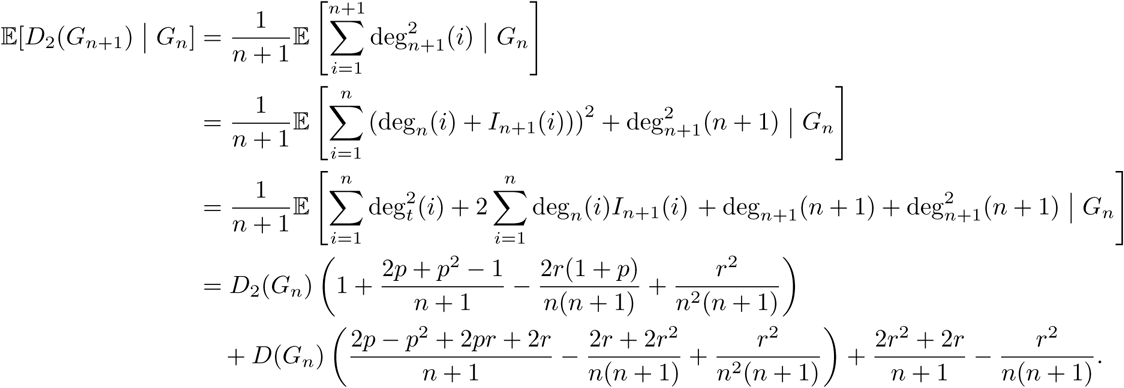

Then from the law of total expectation:

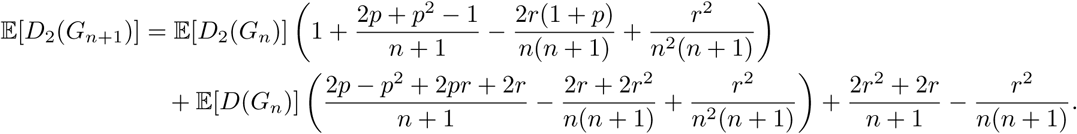

#### 3. Recurrence for *S*_2_(*G*_*n*_)

The recurrence for *S*_2_(*G*_*n*_) is straightforward from the following *deterministic* relation for every graph:

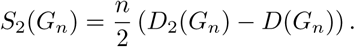

Alternatively, it can be computed from the following RELATION using similar methods as before:

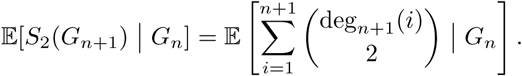

#### 4. Recurrence for *C*_3_(*G*_*n*_)

Finally, we find the expected number of triangles in the following way. Let us denote by *e*_*t*_(*A, B*) the number of edges with one endpoint in *A* and other in *B*, and by *e*_*t*_(*A*) the number of edges with both edges in *A* for some fixed *A, B* ⊆ *V* (*G*_*t*_), at time *t*.

For brevity, let us also introduce the following notations.

- *X*_1_(*G*_*n*_): = *e*_*n*_(𝒩_*n*_(parent(*n* + 1))) – the number of edges within (open) neighborhood of the parent of *n* +1 in *G*_*n*_,
- *X*_2_(*G*_*n*_): = *e*_*n*_ (𝒩_*n*_(parent(*n* + 1)), *V* (*G*_*n*_) 𝒩_*n*_(parent(*n* + 1))) – the number of edges between (open) neighborhood of the parent of *n* + 1 and other vertices in *G*_*n*_,
- *X*_3_(*G*_*n*_): = *e*_*n*_ (*V* (*G*_*n*_) \ 𝒩_*n*_(parent(*n* + 1))) – the number of edges between vertices not connected to the parent of *n* + 1 in *G*_*n*_.

It can be easily verified that

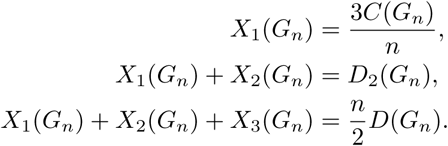

Therefore,

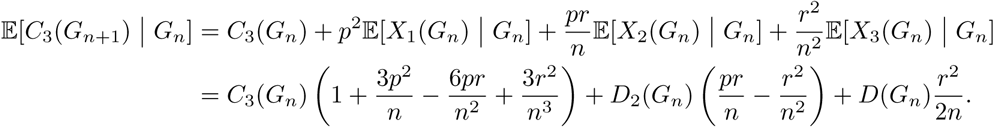

We can now apply the law of total expectation to get the final result.

## Footnotes

2 The subscript *k* in *G*_*k*_ can also be interpreted as time instant *k*.

3 https://thebiogrid.org

